# Perceptual warping exposes categorical representations for speech in human brainstem responses

**DOI:** 10.1101/2022.07.13.499914

**Authors:** Jared A. Carter, Gavin M. Bidelman

## Abstract

The brain transforms continuous acoustic events into discrete category representations to downsample the speech signal for our perceptual-cognitive systems. Such phonetic categories are highly malleable and heir percepts can change depending on surrounding stimulus context. Previous work suggests these acoustic-phonetic mapping and perceptual warping of speech emerge in the brain no earlier than auditory cortex. Here, we examined whether these auditory-category phenomena inherent to speech perception occur even earlier in the human brain, at the level of auditory brainstem. We recorded speech-evoked frequency following responses (FFRs) during a task designed to induce more/less warping of listeners’ perceptual categories depending on stimulus presentation order of a speech continuum (random, forward, backward directions). We used a novel clustered stimulus paradigm to rapidly record the high trial counts needed for FFRs concurrent with active behavioral tasks. We found serial stimulus order caused perceptual shifts (hysteresis) near listeners’ category boundary confirming identical speech tokens are perceived differentially depending on stimulus context. Critically, we further show neural FFRs during active (but not passive) listening are enhanced for prototypical vs. category-ambiguous tokens and are biased in the direction of listeners’ phonetic label even for acoustically-identical speech stimuli. Our data expose FFRs carry category-level information and suggest top-down processing actively shapes the neural encoding and categorization of speech at subcortical levels. These findings suggest the acoustic-phonetic mapping and perceptual warping in speech perception occur surprisingly early along the auditory neuroaxis, which might aid understanding by reducing ambiguity inherent to the speech signal.

## 1. INTRODUCTION

To effectively utilize speech, individuals must convert continuous stimuli in the external world to phonetic category units (Goldstone and Hendrickson, 2010). In continuous speech, the precise acoustic characteristics of phonemes vary depending on the speaker (e.g., sex, accent) (Sumner, 2011), surrounding coarticulation (Beddor et al., 2002), and background noise (Bidelman, 2016; Billings et al., 2009; Carter and Bidelman, 2021). Categorization allows this variation to exist without hindering the utility of speech as a mode of communication. One open question in categorization is whether its driving force lies in neurophysiological constraints of the sensory system (i.e., bottom-up coding of sound) (Kuhl, 1986; Kuhl and Miller, 1975) or if higher-order language and memory regions modulate category speech percepts in a top-down manner (Bidelman et al., 2021; Carter and Bidelman, 2021; Ganong III and Zatorre, 1980; Kuhl, 1986; Kuhl and Miller, 1975). If top-down modulations of early speech representations do occur, then how far down the auditory system are these perceptual influences exerted?

Typically, when assessing categorization, signals are presented to listeners who are asked to identify the sound as a member of a set of discrete categories. Their behavioral responses can be represented as a psychometric function, which can be quantified by its slope and its categorical boundary. A steeper slope indicates the perceptual change from one category to the next happens more rapidly than if the slope was shallower and thus indexes the strength of categorical hearing across the continuum (Bidelman, 2015a; Strouse et al., 1998; Xu et al., 2006a). The categorical boundary indicates the point at which the psychometric function crosses 50% identification, marking the stimulus location where the category shifts from one percept to another (Altmann et al., 2014; Ganong III and Zatorre, 1980). Additionally, one can measure how rapidly a listener labels each token via reaction time (RT). RTs demonstrate the speed of processing, which increases (i.e., slows down) during more ambiguous or degraded tokens and decreases (i.e., speeds up) during more prototypical tokens, yielding an inverted U shape when plotting RTs across the continua (Pisoni & Tash, 1974).

The categorical perception of speech is usually assessed by randomizing the presentation of stimuli from a graded acoustic continuum. When presenting stimuli in sequential order (e.g., high-to-low first formant frequency [F1]), rather than a random order, the categorical boundary is modified due to short-term sequencing effects (Diehl et al., 1978; Healy and Repp, 1982). Such perceptual shifts may reflect a perseveration of the prior perception (i.e., hysteresis) or changing perception to the other category earlier than anticipated (i.e., enhanced contrast) (Tuller et al., 1994). These types of dynamics in perception suggest the brain’s ongoing sorting of incoming acoustics into categorical phonetic representations is actively modulated in a top-down manner. When viewed through the lens of nonlinear dynamic systems, this process can be described as a shifting of the perceptual space to accommodate variability within categories (Tuller et al., 1994). Such warpings in perceptual space are likely driven by prefrontal (i.e., memory) brain regions that track ongoing stimulus history and adjust current percepts according to listeners’ expectations and perceptual biases (Carter et al., 2022; Hansen et al., 2006). We do not yet know how far down the auditory system this top-down modulation of speech representation continues, however. While fronto-temporal pathways drive auditory stimulus encoding in cortex, the corticofugal system (i.e., cortico-collicular efferent pathways) can also modulate responses in the auditory brainstem by fine-tuning sound representations according to listening demands (Suga, 2008; Suga et al., 2000). Additionally, corticofugal fibers enhance speech processing prior to its arrival in cortex through attention-dependent gain control (Lai et al., 2022a; Price and Bidelman, 2021). This makes the corticofugal system a prime candidate for tuning speech representations and possibly building nascent acoustic-phonetic structure at *subcortical* levels.

The frequency-following response (FFR) has been used as a window to characterize early, subcortical sound encoding along the auditory system. The FFR is a scalp-recorded potential evoked by sustained stimuli (such as speech) occurring ∼7-10 milliseconds after stimulus onset with putative source(s) in the auditory brainstem (i.e., inferior colliculus) (Bidelman, 2018b; Gardi et al., 1979; Langner and Schreiner, 1988; Smith et al., 1975; Sohmer et al., 1977), and not cochlear origin (Skoe and Kraus, 2010). Early work in animal models localized the FFR to several subcortical auditory nuclei including cochlear nucleus (CN), inferior colliculus (IC), and medial geniculate body (MGB) (Dunlop et al., 1965; Oatman and Anderson, 1980; Sohmer et al., 1977). While most previous work has shown a subcortical origin of the FFR, recent neuroimaging studies have revealed cortical contributions to the response at low (∼100 Hz) speech frequencies when recorded via magnetoencephalography (MEG) (Coffey et al., 2016b). However, EEG work has convincingly demonstrated that subcortical structures (i.e., midbrain and even auditory nerve) provide the largest contribution to the scalp-recorded FFR_EEG_ for most of the frequency bandwidth of speech (Bidelman, 2018b; Bidelman and Momtaz, 2021). The FFR phase-locks with the time-varying, spectro-temporal features of complex sounds including fundamental frequency (F0) and harmonics (Galbraith et al., 1995), as well as the first few formant frequencies up to its phase locking limits (∼1200 Hz) (Aiken and Picton, 2008; Krishnan, 2002; Skoe and Kraus, 2010). Given its unique time-frequency signature within the EEG, FFRs have been used to characterize subcortical processing of speech (Bidelman and Powers, 2018; Bidelman and Momtaz, 2021; Bidelman et al., 2013a; Galbraith et al., 1995; Johnson et al., 2005; Musacchia et al., 2008; Russo et al., 2004; Skoe and Kraus, 2010) and musical sounds (Bidelman, 2013; Bones et al., 2014; Mankel and Bidelman, 2018), as well as track changes in neural encoding across the lifespan (Anderson et al., 2012; Bidelman et al., 2019; Bidelman et al., 2014a; Liu et al., 2018). Of interest for this study is the use of FFRs in understanding the brain’s earliest neural representations for speech and its sensitivity to specific phonetic features found in a listeners’ native language (cf. categories) (Krishnan et al., 2010; Krishnan et al., 2009).

To date, categorical representations have not been observed in brainstem FFRs, which, despite their ability to faithfully encode speech stimulus properties (e.g., formants), do show strong evidence of category structure. This is despite concomitant category representations observed in the same listeners’ cortical evoked potentials (Bidelman et al., 2013a). However, categorization in most neuroimaging studies is tested under active tasks; the passive listening task used in subcortical studies is known to reduce category coding in neural responses (Alho et al., 2016; Bidelman and Walker, 2017). Nevertheless, some evidence exists showing the possibility of category representation in subcortical structures. In guinea pig, auditory brainstem responses evoked by noise bursts separate in a nonlinear fashion (indicative of categorical coding) based on the gap duration between noise bursts (Burghard et al., 2019). Studies that compared listeners fluent in tonal (e.g., Chinese) vs. non-tonal (e.g., English) languages show that the former tend to have stronger pitch representation in subcortical responses, but only for pitches that match native pitch contours in their language (Krishnan et al., 2010; Krishnan et al., 2009; Xu et al., 2006a). However, this effect does not carry over to similar acoustic analogues of the pitch changes that are not found in the native tone space (Xu et al., 2006b). Such findings suggest the presence of linguistically-relevant (categorical-like) information in the brainstem, but itself does not indicate the active process of categorization is occurring de novo in brainstem, *per se*. Such findings could be explained by long term, experience-dependent plasticity (Krishnan et al., 2012). This evidence is further bolstered by findings of categorization-training studies that show once individuals learn to identify novel speech stimuli their FFR are enhanced relative to more novice listening states (Cheng et al., 2021; Reetzke et al., 2018).

A possible mechanism that would enable FFRs to show real-time category representations is attention/behaviorally-dependent control of the corticofugal pathway. Attention heavily modulates responses from cortical structures (Bidelman and Walker, 2017; Harris et al., 2012; Hillyard et al., 1973; Zhang et al., 2014). It is perhaps expected then that categorical representations in the (cortical) event-related potentials (ERPs) are only observed under states of attentional load and active speech labeling tasks (Alho et al., 2016; Bidelman and Walker, 2017; Carter, 2018). Literature on attentional effects in human brainstem responses is mixed with some suggesting attentional enhancement of FFRs (Galbraith et al., 1998; Hartmann and Weisz, 2019; Price and Bidelman, 2021) while others finding little to no effect of attention on the FFR (Aiken and Picton, 2008; Dunlop et al., 1965; Galbraith and Kane, 1993; Varghese et al., 2015). If attention does influence the brainstem FFR, then actively categorizing speech during behavioral tasks should yield measurable changes the response. Moreover, stimulus order effects in the FFR would provide new evidence that subcortical speech representations are not only influenced by local stimulus history but are indeed tuned by nonlinear perceptual dynamics as observed at a cortical level (Carter et al., 2022).

The current study aimed to evaluate (1) if speech representations, as indexed by brainstem FFRs, show evidence of categorical representation or are strictly sensory-acoustic depictions of the speech signal; (2) whether attention and the process of categorization actively modulate speech-FFRs; (3) the effects of nonlinear dynamics (i.e., perceptual warping) on brainstem representations for speech. To this end, we measured speech-FFRs while listeners performed a rapid phoneme identification task where tokens along an identical categorical continuum were presented in random vs. serial (forward or backward) order. This design allowed us to induce more/less perceptual warping to bias listeners’ categorical hearing. Serial order warps the perceptual space and corresponding *cortical* acoustic-phonetic representations for speech (Carter et al., 2022). Here, we determined if *subcortical* FFRs similarly carry category-level information that also changes with listeners’ ongoing speech percept. We measured F0 and F1 attributes from FFRs to quantify “voice pitch” and “formant timbre”-related coding in neural responses. We first confirmed our novel paradigm shifted individual’s perceptual categorical boundary measured behaviorally and thus successfully warped (biased) listeners’ percept. If brainstem speech coding is sensitive to categorization, we hypothesized FFRs would show larger F1 amplitudes in sequential vs. random presentation orders. We also anticipated relationship between neural and behavioral measures if the FFR is indeed modulated by listeners’ ongoing categorical percept. Our data reveal that category-level features of speech are actively coded in FFRs, suggesting the acoustic-phonetic mapping of speech occurs more peripheral in the auditory system than previously thought.

## 2. MATERIALS & METHODS

### 2.1 Participants

The sample included N=16 young participants (24.2 ± 4.4 years; 5 females) averaging 16.9 ± 3.2 years of education; n=9 of these listeners also participated in Carter et al. (2022). All spoke American English, had normal hearing (air conduction thresholds ≤20 dB HL; 250–8000 Hz), minimal musical training (≤3 years; average = 0.9 ± 1.2 years), and were mostly right-handed (mean = 78% ± 29% laterality) (Oldfield, 1971). Each gave written informed consent in compliance with a protocol approved by the University of Memphis IRB.

### 2.2 Stimuli & task

We used a 7-token vowel continuum from /u/ to / /. Each 100 ms token had a fundamental frequency of 150 Hz to avoid cortical contributions to the FFR which are restricted to low (< 100-120 Hz) stimulus frequencies (Bidelman, 2018b; Brugge et al., 2009). Adjacent tokens were separated by equidistant steps in first formant (F1) frequency spanning from 430 (/u/) to 730 Hz (/ /). We selected vowels over consonant-vowel (CV) syllables because our prior work showed vowels were more prone to nonlinear perceptual effects than stop consonants (Carter et al., 2022). We delivered stimuli binaurally through insert earphones (ER-2) at 80 dB SPL with electrical shielding to prevent stimulus electromagnetic artifact from contaminating neural responses (Campbell et al., 2012; Price and Bidelman, 2021). Sound presentation was controlled by MATLAB coupled to a TDT RZ6 signal processor (Tucker-Davis Technologies, Alachua, FL).

FFRs are sub-microvolt signals that require minimally 1000 trials for response detection (Bidelman, 2018a). To use our categorization paradigm while simultaneously recording FFRs, we employed a modified version of the clustered interstimulus interval (ISI) presentation paradigm as described in Bidelman (2015c). This grouped stimuli in blocks containing rapid bursts of the same token (20 repetitions; ISI = 10 ms) within a short train. After each train, the participant selected the phoneme they perceived in the group with a binary keyboard response (“u” or “a”), after which the ISI was slowed (ISI = 400 ms) before the next grouping. The clustered ISI sequence was then repeated to achieve the appropriate token counts to detect the FFR (x1000 presentations per token per condition) and sufficient behavioral responses (x50 per token) (Figure 1).

**Figure 1.**
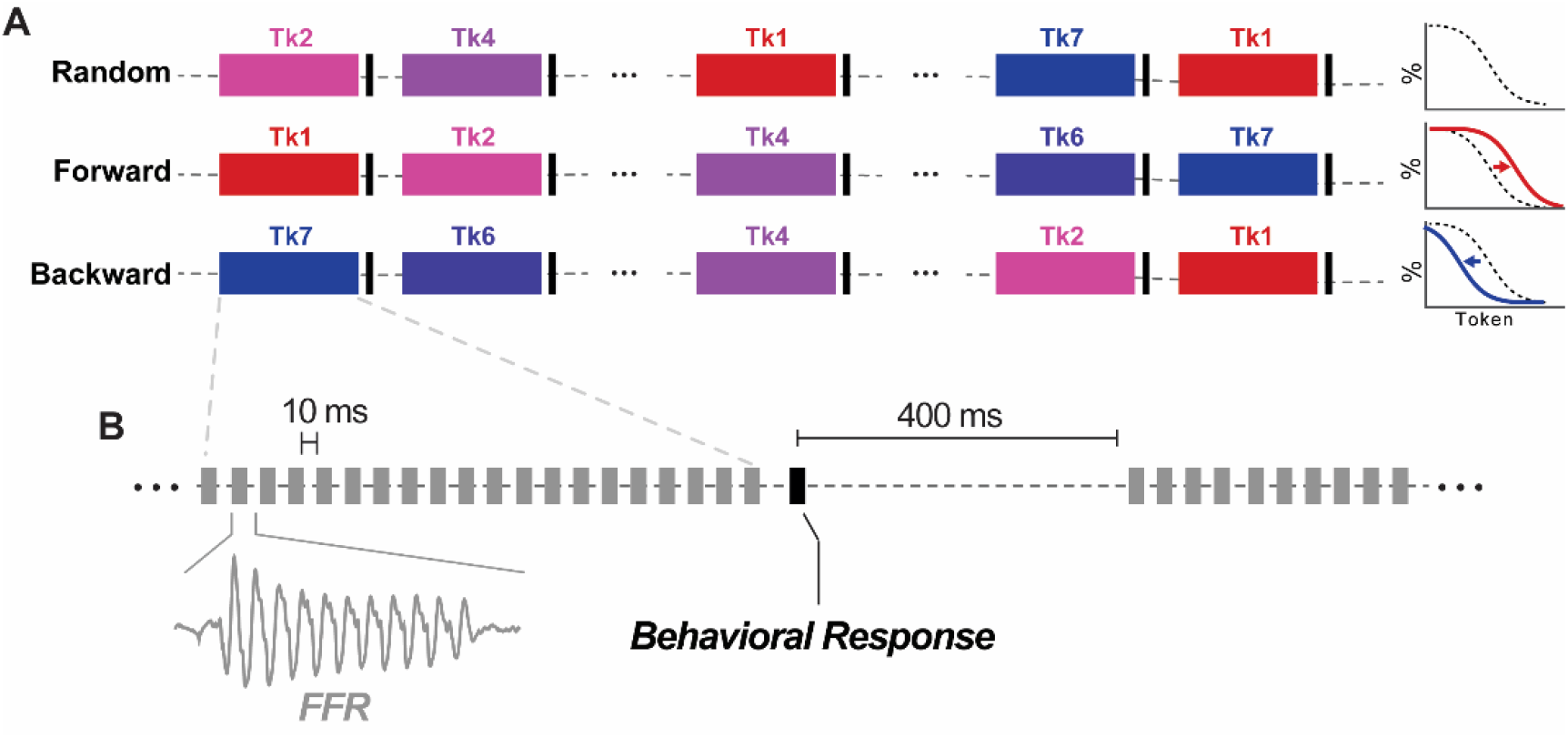
**(A)** Schematic of the presentation orders and examples of changes to the psychometric function the presentation order could create. **(B)** Schematic of the stimulus clustering paradigm for recording FFRs during active behavioral tasks [modified from Bidelman (2015c)]. Speech tokens were presented rapidly in blocks of twenty (10 ms ISI) to evoke the FFR. At the end of the block, stimuli were paused and the listener categorized the sound as /u/ or / /. Following the behavioral response, a 400 ms pause occurred and the next block was presented. The clustered ISI sequence is then repeated to achieve the appropriate token counts for the FFR (x1000 presentations per token per condition) and sufficient behavioral responses (x50 per token).

There were three active conditions based on how tokens were sequenced: (1) random presentation, and two sequential orderings presented serially between continuum endpoints and F1 frequencies (2) forward /u/ to / /, 430 Hz to 730 Hz, and (3) backward / / to /u/, 730 to 430 Hz). Forward and backward directions on such a continuum were expected to produce perceptual warpings (i.e., hysteresis) (Tuller et al., 1994). An additional passive condition in which the stimuli were presented in a random order while the listeners watched a captioned film of their choice (but ignored the vowel stimuli) was used to test for attention effects on the FFR. The conditions were pseudo-randomly assigned using a Latin Square counterbalance (Bradley, 1958). In a subset of listeners (*n*=5), we measured the noise floor of our FFR recording setup to rule out electromagnetic contamination of the neurophysiological recordings. This used an identical setup to the passive block only with the earphone removed from listeners’ ear canal thereby recording only “neural noise” (Price and Bidelman, 2021).

### 2.3 EEG recording procedures

Neuroelectric activity was recorded between Ag/AgCl electrodes placed on the high forehead scalp (∼Fz) referenced to linked mastoids (M1/M2) (with a mid-forehead electrode as ground), as is standard for recording brainstem FFRs (Billings et al., 2019; Coffey et al., 2016a; Gockel et al., 2013; Skoe and Kraus, 2010). This montage is used to obtain responses from the vertically oriented dipoles in the brainstem (Bidelman, 2015b; Chandrasekaran and Kraus, 2010). Interelectrode impedances were kept ≤6 kΩ. EEGs were digitized at 10 kHz. Responses were epoched (−5 – 105 window), artifact rejected (set to reject the highest 5% of response amplitudes in each run) and averaged to derive FFRs for each vowel stimulus. Responses were then band-passed filtered (130 – 2000 Hz) for visualization and quantification. This passband effectively attenuates cortical activity of the EEG while maintaining the high spectral resolution of the speech-FFR including the voice F0 and its harmonics captured in the response (Bidelman et al., 2013a; Musacchia et al., 2008).

### 2.4 Behavioral data analysis

#### 2.4.1 Psychometric Function Analyses

Identification scores were fit with sigmoid P = 1/[1 + e^−β1(x−β0)^], where P is the proportion of trials identified as a given vowel, x is the step number along the continuum, and β0 and β1 are the location and slope of the logistic fit estimated using non-linear least-squares regression (Bidelman and Walker, 2019; Bidelman et al., 2014b). Leftward/rightward shifts in β0 location for the sequential vs. random stimulus orderings would reveal changes in the perceptual boundary characteristic of perceptual nonlinearity (Tuller et al., 1994). RTs greater than 2500 ms were considered outliers (e.g., attention lapses) and were excluded from analysis (reject trials: 208; 1.23% across all conditions/subjects/tokens) (Bidelman et al., 2013a; Bidelman and Walker, 2019). We included RTs ≤250 ms, as we expected the task to induce faster RTs given a quasi-priming (anticipation) effect of the stimulus sequencing where the listener might decide a percept during the ongoing token train.

#### 2.4.2 Classifying Response Patterns

To classify participants based on their different listening strategies (hysteresis/enhanced contrast/critical boundary), we calculated the standard deviation of the categorical boundary across all listeners in the random condition. We then considered whether each individual’s categorical boundary fell within a 2SD around the group mean for the forward/backward. Participants whose boundary fell within 2SD were categorized as critical boundary listener. If their categorical boundary occurred before this window (e.g., Tk3 for backward; Tk5 for forward), we categorized them as hysteresis listener. If instead their categorical boundary happened after this window (e.g., Tk5 for backward; Tk3 for forward), we categorized them as an enhanced contrast listener (Carter et al., 2022).

### 2.5 Electrophysiological data analysis

#### 2.5.1 FFR analysis

FFR analyses were conducted using automated routines coded in MATLAB. We computed the Fast Fourier Transform (FFT) of each FFR to assess spectral content in each waveform. We then measured the F0 and F1 of the spectra as the maximal FFT amplitude in a window ±50 Hz around the nominal stimulus F0 and F1 frequencies. As voice pitch (F0) was identical across our stimuli, we expected FFR F0 to remain invariant across tokens and sequence orders. In contrast, we expected differences in FFR F1 amplitudes where the stimuli are systematically changed to create the categorical continuum. We compared the FFT amplitudes of F0 and F1 of the same stimulus in different presentation conditions. Although not indicative of categorization, we also expected differences in F1 frequency across tokens since the FFR closely tracks changes in stimulus acoustics and we changed F1 frequency by the stimulus design [see Fig. 2d; Bidelman et al. (2013a)].

**Figure 2.**
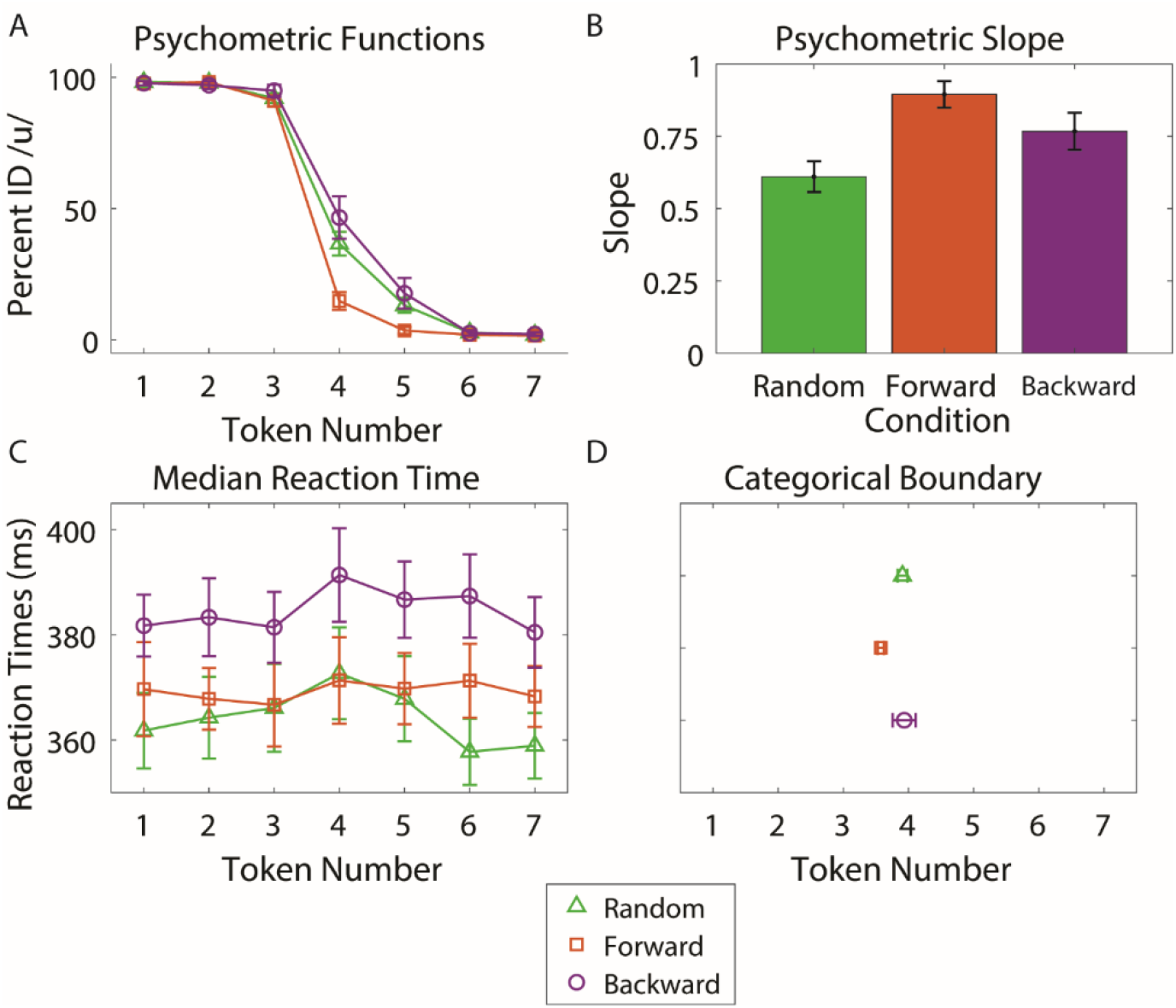
Group level behavioral categorization. **(A)** Perceptual psychometric functions for phoneme identification when continuum tokens are presented in random vs. serial (forward: /u/→/ / vs. backward: / /→/u/) order. **(B)** Psychometric function slope was steeper for serial (forward and backward) compared to random presentation order. **(C)** Reaction times for speech identification. Backward presentation led to slower RTs than random and forward presentations. Additionally, there was no token difference in RTs. **(D)** Boundary location did not vary at the group level (cf. individual differences; Fig. 3). Errorbars = =±1 s.e.m.

#### 2.5.2 Neural adaptation

To determine if neural adaptation occurred given the repetitive nature of our stimulus trains, we compared the F0 amplitude of the first and the last token in each train (for each continuum token: Tk1-Tk7). We compared the amplitude of F0 between the first and the last tokens to determine if the neural responsiveness decreased with the rapid presentation of the stimuli within the trains. This would indicate the fast repetition of speech stimuli caused adaptation of neural responses (Pérez-González and Malmierca, 2014). Adaptation might inadvertently account for differential amplitude changes with stimulus presentation order and confound our interpretations of hysteresis and categorical representations in the FFR.

#### 2.5.3 Response-to-response correlations

To determine if stimulus ordering and thus perceptual warping biased listeners’ speech-FFRs we measured response-to-response correlations between FFRs to the ambiguous token (Tk4) and the two endpoints (Tk1, Tk5) (cf. Yellamsetty and Bidelman, 2019). For each listener, we cross-correlated their time waveform to Tk4 for each serial order (forward, backward) with their time waveforms to both prototypical endpoints (Tk1 and Tk7 in the random condition). Waveforms were allowed to shift up to ±10 ms relative to one another to account for differences in delays (Galbraith and Brown, 1990). This resulted in four correlation coefficients per listener, reflecting the degree to which the FFR to the otherwise identical speech sound (Tk4) mirrored each of the two categories. We reasoned that if the ambiguous token is more like one of the prototypical tokens than the other as a function of direction, it would indicate that the encoding of the signal was modulated by the perceptual warping induced by recent stimulus history (Yellamsetty and Bidelman, 2019).

#### 2.5.4 Neural decoding of speech categories from single-trial FFRs

We used Gaussian kernel, linear support vector machine (SVM) classifiers to determine if the categorical identity of speech stimuli could be decoded via FFRs (e.g., Lai et al., 2022b; Xie et al., 2019; Yi et al., 2017). We used log-transformed F0/F1 amplitudes as the input features for SVM decoding. Amplitude measures were selected, as opposed to F0/F1 frequencies, as the latter would be trivial in separating FFRs since they closely follow the low-frequency pitch and formant cues of speech (see Fig. 2 of Bidelman et al., 2013b). Two binary classifications were considered: models assessing the classification of (i) FFRs elicited by the two category prototypes (i.e., Tk1=/u/ vs. Tk7=/ /) and (ii) FFRs elicited by the otherwise ambiguous Tk4 in forward vs. backward serial directions. The first model was used largely as a control analysis since we expected robust separability of FFRs to differing vowel tokens and thus near ceiling classifier performance. In these analyses, for example, the classifier attempted to predict the FFR response on a given trial (F0amp_n_) as being evoked by either an /u/ or / / stimulus. Of more interest was the second model, which tested whether FFRs to a category-ambiguous speech sounds were warped based on listeners’ trial-to-trial phonetic hearing.

For each classifier, we extracted F0 and F1 amplitudes from *single-trial* FFR spectra, resulting in N=27000 observations per condition. We randomly split the data into training (80%) and test (20%) sets (Mahmud et al., 2021). We then trained an SVM via the *fitckernel* function in MATLAB using the default box constraint (*C=1*) and regularization (λ=1/*n =* 4.63e-05) tuning parameters, where *C* = 1/(λ*n*). This algorithm maps data from a low- to high-dimensional space, then fits a linear model in the high-dimensional space by minimizing the regularized objective function. We used 5-fold cross validation to prevent model overfitting (Mahmud et al., 2021). In this procedure, the dataset is partitioned into *k* equal-sized subsamples (folds) containing N=21600 (80% training) and N=5400 (20% testing) observations. For each fold, the SVM learned the support vectors from the training data that optimally segregated the FFR attributes (i.e., F0 and F1 amplitudes) based on the class labels (e.g., Tk1 vs. Tk7 or Tk4_for_ vs. Tk4_back_). This was repeated for each fold. The final classifier performance represented the mean decoding accuracy (i.e., % of matches between predicted and true class labels) averaged across folds. Classification performance was then assessed using conventional classifier metrics [i.e., accuracy, d-prime, receiver operating characteristic curve (ROC), confusion matrices].

### 2.6 Statistical analysis

We used one-way mixed model ANOVAs (PROC GLIMMIX, SAS® 9.4; SAS Institute, Inc.) to analyze the psychometric data, with a fixed effect for presentation condition (3 levels: random, forward, and backward), and a random effect for subjects. RTs and FFR data (i.e., F0 and F1 frequency and amplitudes) were analyzed using a two-way, mixed model ANOVAs (subjects = random factor) with fixed effects of condition (3 levels: random, forward, backward; 4^th^ level for FFR: passive) and token (7 levels). We normalized the heavily bimodal distribution of the correlation data by taking the absolute value of the difference of the individual’s correlation value and the mean of all correlations (i.e., abs(X - mean(X)).

We used orthogonal quadratic trend contrasts on F0 and F1 amplitude measures to test for the characteristic U-shape pattern inherent to categorical responses (Pisoni, 1973). These *a priori* contrasts (coefficients = 5, 0, −3, −4, −3, 0, 5) assessed whether FFR amplitudes to token prototypes were larger (or smaller) than ambiguous tokens near the continuum’s midpoint (Carter and Bidelman, 2021; Mankel et al., 2020) and therefore differentiated speech sounds with strong vs. weak category percepts. We anticipated that if category-level information is encoded in brainstem responses, a similar quadratic trend would arise.

We conducted general linear mixed effects (GLME) regression models (*fitglme* in MATLAB) to assess whether a linear combination of the neural measures (i.e., F0/F1 frequencies and amplitudes) predicted behavior [e.g., behav ∼ FFR_F0amp_+ FFR_F0freq_ + FFR_F1amp_+ FFR_F1freq_ + (1|sub)]. Subjects served as a random factor in these models. Separate GLMEs were run for each behavioral metric (i.e., slope; boundary; RTs). Responses across the 3 orders were pooled for data reduction.

## 3. RESULTS

### 3.1 Behavioral data

Listeners perceived the vowels categorically in all presentation orderings as seen in **Figure 2**. Slopes varied with presentation order (*F_2,30_*= 11.21, p = 0.0002). The random condition was significantly shallower than both the forward (*p* = 0.0001) and backward (*p* = 0.0367) conditions. The location of the categorical boundary only showed marginal shifts with presentation order at the *group level* (*F_2,30_* = 3.14, *p* = 0.0576). These findings are consistent with notions that categorical speech percepts are stronger when stimuli are presented in a sequential compared to random order (Carter et al., 2022).

RTs also varied with presentation order (*F_2,312_* = 18.72, *p* < 0.0001). Categorical decisions were slower for backward versus forward (*p* < 0.0001) and random (*p* < 0.0001) presentations. This finding indicates that the backward condition slowed processing speed in categorization. However, there was no difference in the RTs between ambiguous and prototypical tokens (*F_6,312_* = 1.41, *p* = 0.21). This suggests that under our clustered stimulus paradigm, listeners may have decided the category while the stimulus train was still ongoing.

While the group level categorical boundary was largely stagnate, individual-level data showed stark differences as a function of presentation order (**Fig. 3**). In the backward condition, most listeners retained a critical boundary (*n* = 11), some showed enhanced contrast (*n* = 4), and one showed hysteresis (*n* = 1). In the forward condition, even more listeners demonstrated a critical boundary (*n* = 13) with some listeners having enhanced contrast (*n* = 3).

**Figure 3.**
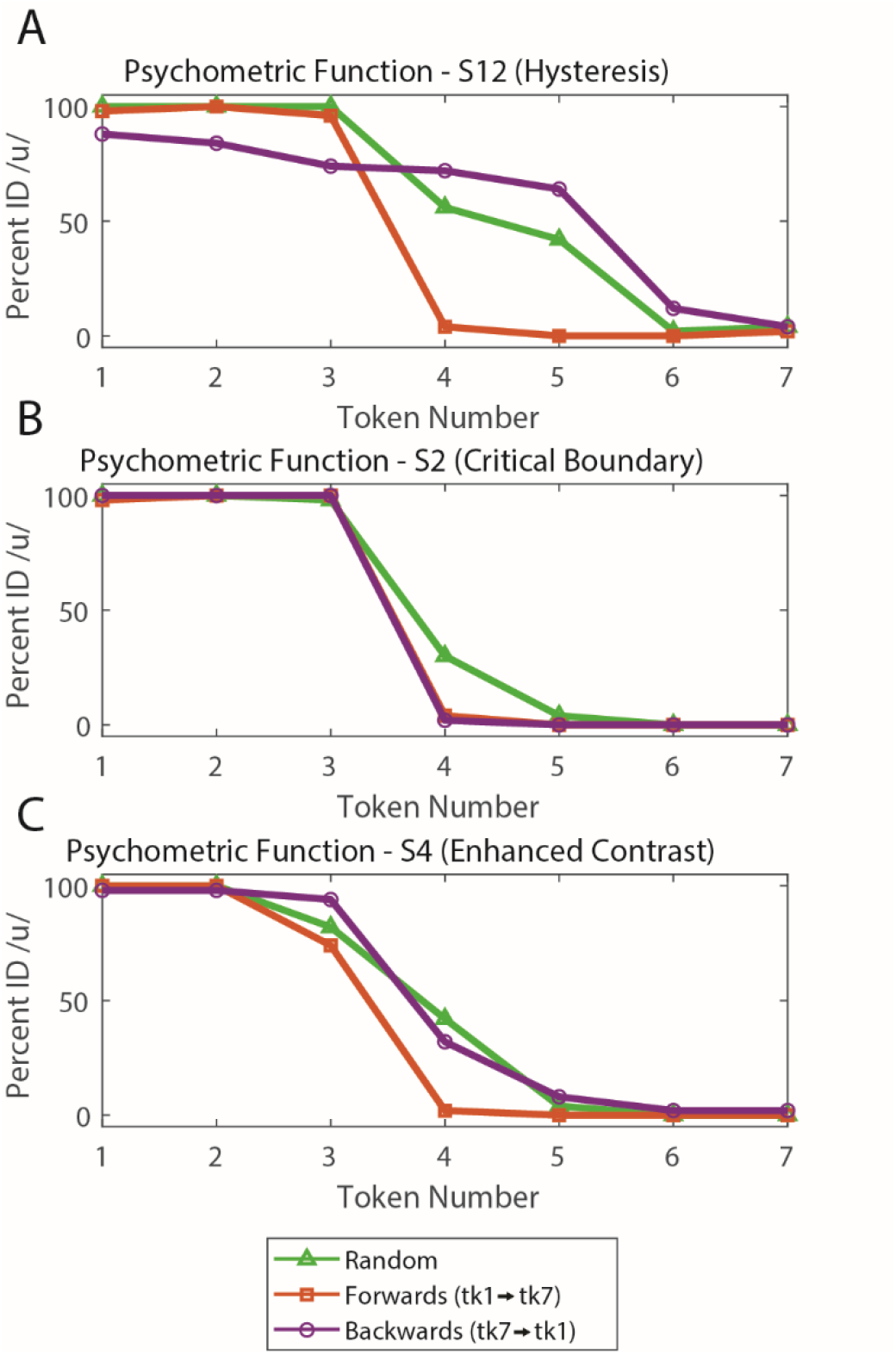
Individual level psychometric functions. Representative subjects who showed **(A)** hysteresis **(B)** critical boundary and **(C)** enhanced contrast perceptual response patterns.

### 3.2 Electrophysiological data

**Figure 4** and **Figure 5** show time-domain FFR waveforms for select conditions and tokens. These waveforms were analyzed in the frequency domain to determine differences in F0 and F1 frequency and amplitude.

**Figure 4.**
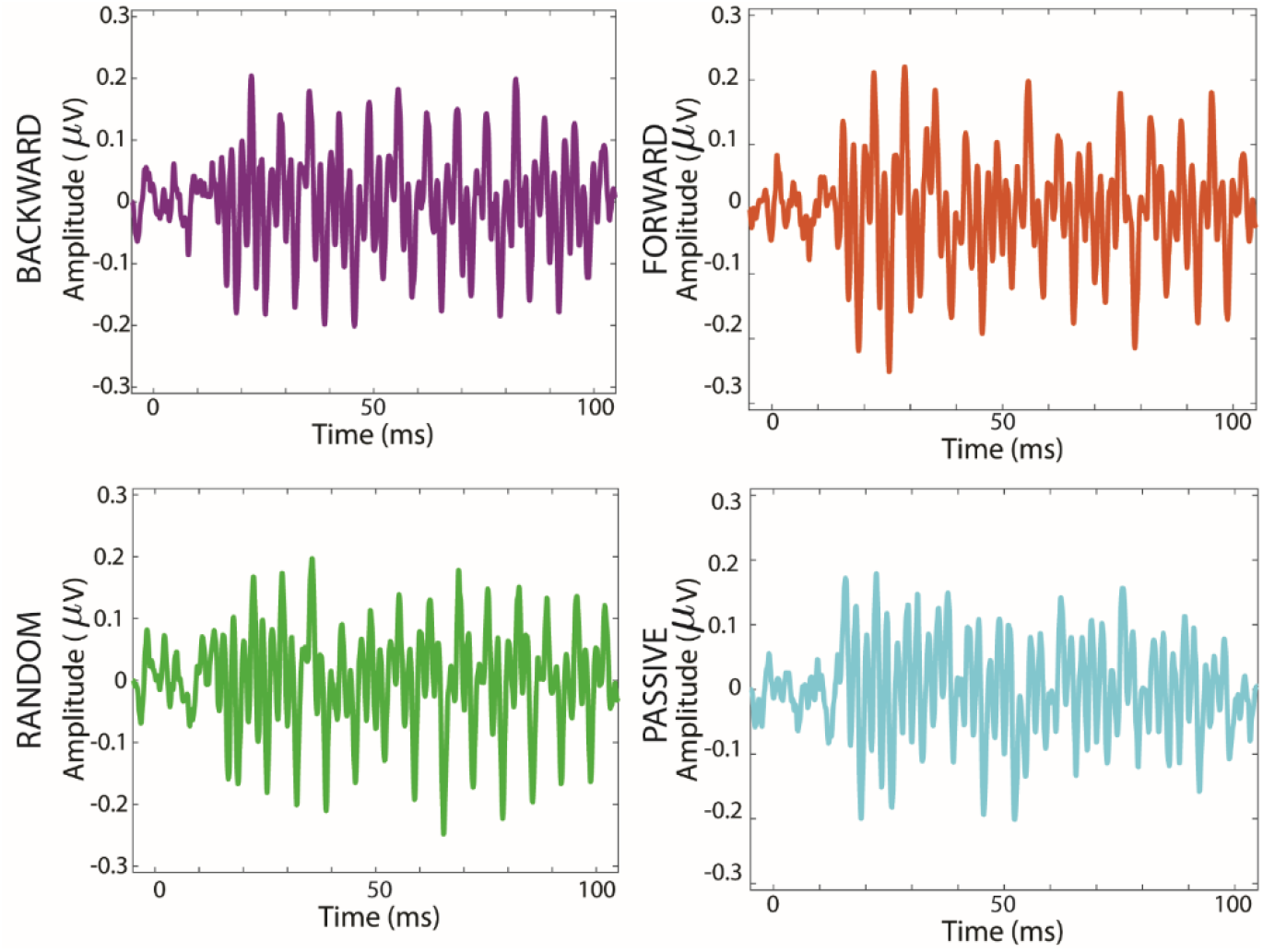
FFR time domain waveforms (Tk1) contrasting stimulus presentation orders and attentional state (i.e., active vs. passive listening).

**Figure 5.**
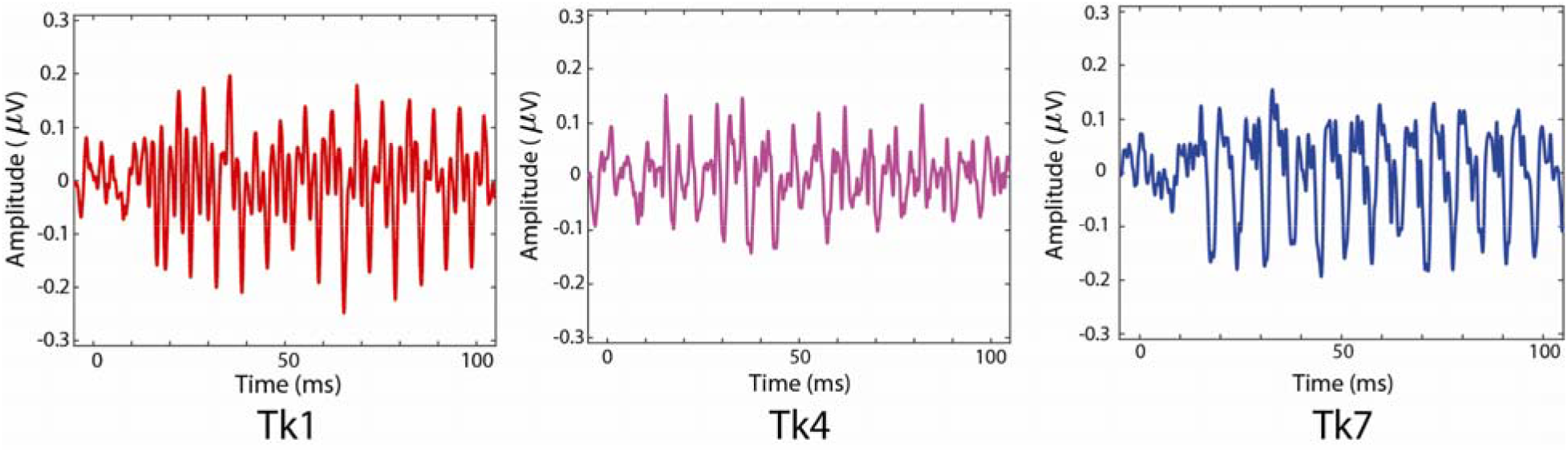
FFR time domain waveforms comparing responses to mid- and endpoint tokens (Tk1, Tk4, Tk7) for the random condition.

**Figure 6** shows FFR spectra in response to Tk1 across stimulus orderings (random, forward, backward) and attention conditions. **Figure 7** shows F0 and F1 measures more clearly. We found that F0 amplitude differed as a function of condition (*F_3,423_*= 7.91, *p* < 0.0001) and token (*F_6,423_* = 5.86, *p* < 0.0001). Post-hoc testing revealed the passive F0 amplitudes were smaller than all three active listening conditions (backward, *p* = 0.0004; forward, *p* = 0.0042; random, *p* = 0.0001). The token effect was attributed to smaller F0 amplitudes in response to /u/ tokens (Tks 1-3) compared to / / tokens (Tks 5-7) (*p* < 0.0001). Conversely, F1 amplitude did not differ as a function of condition (*F_3,423_* = 0.31, *p* = 0.8189), but did as a function of token (*F_6,405_* = 131.79, *p* < 0.0001). These results indicate (expectedly) the FFR is sensitive to the acoustic properties of speech across the stimulus continuum. More critical, they indicate subcortical speech representations are enhanced with active attention.

**Figure 6.**
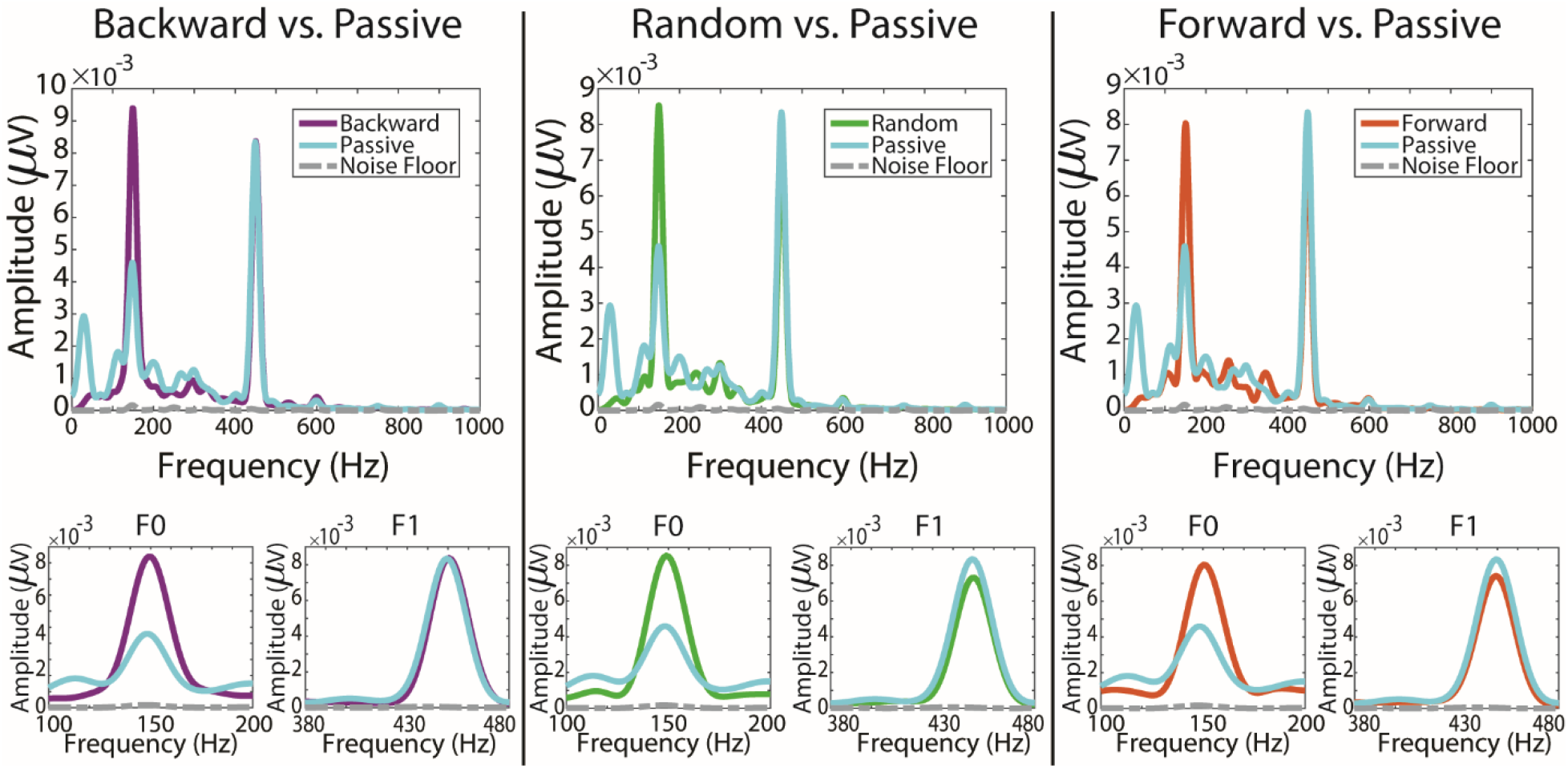
FFR spectra (Tk1) in the backward, random, and forward conditions vs. the passive condition. Insets show F0 and F1 analysis windows.

**Figure 7.**
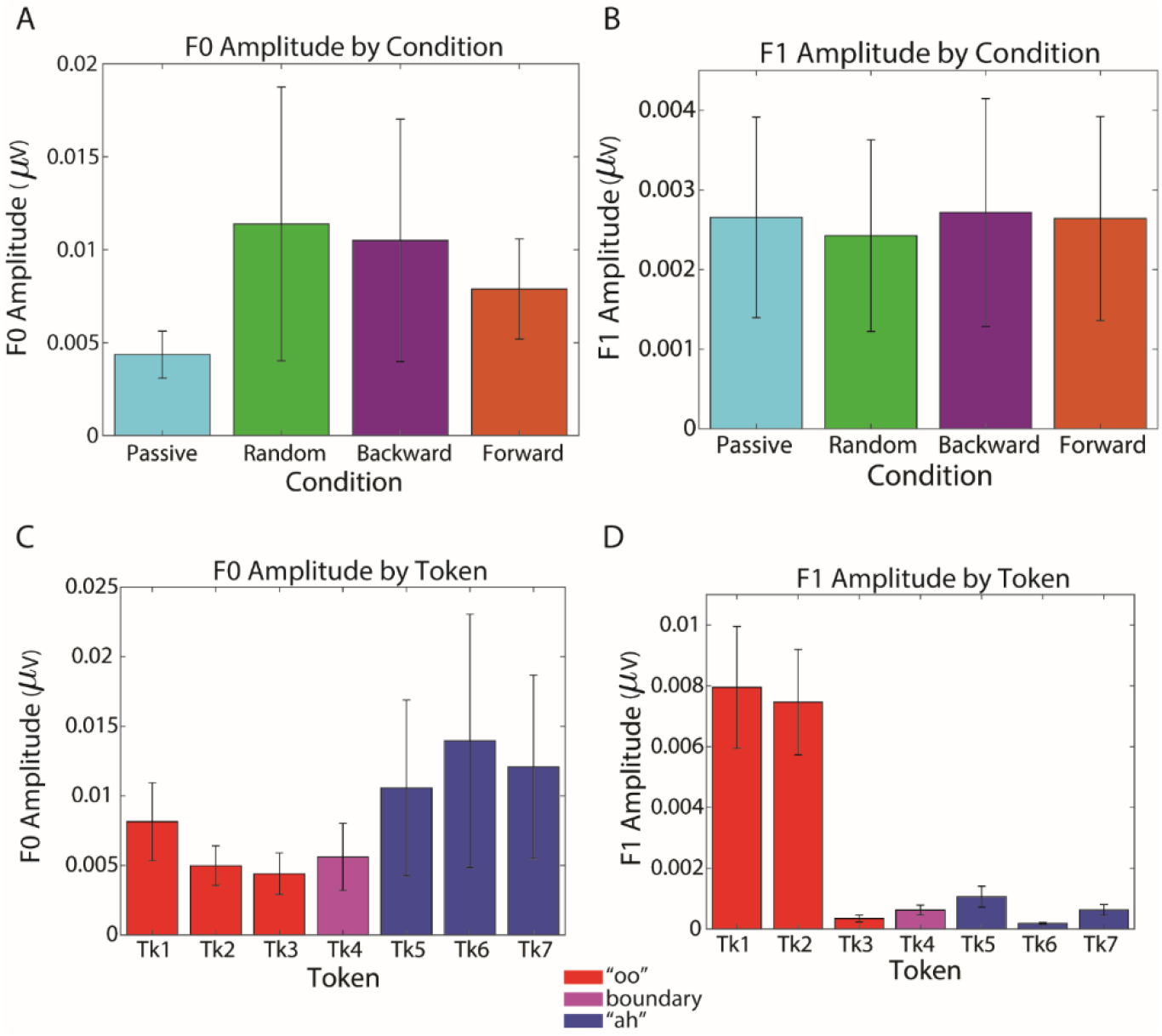
FFR F0 and F1 measures as function of stimulus order and token. **(A)** The F0 amplitude of active conditions (i.e., random, forward, and backward) were greater than the F0 amplitude in the passive condition. **(B)** F1 amplitudes were not affected by stimulus direction nor attention. **(C)** F0 amplitudes (pooling orders) showed a U-shape pattern suggesting categorical coding across speech tokens (Pisoni, 1973). **(D)** F1 amplitudes (pooling orders) were significantly larger for /u/ vs / / ends of the continuum. Errorbars = =±1 s.e.m.

We found FFRs across the categorical continuum displayed a quadratic trend for both F0 (*F_6,423_* = 131.79, *p* < 0.0001) and F1 (*F_6,423_* = 131.79, *p* < 0.0001) measures. Quadratic trends showed a U-shape for the F0 amplitudes in the backward (*p* = 0.0316) and forward (*p* = 0.0149) conditions, but not for the random (*p* = 0.1651) or passive (*p* = 0.5883) conditions. These results indicate that, despite identical F0s in the stimuli, the FFR showed categorical coding of F0 only in sequential presentation orders (which elicited perceptual warping). In contrast, the quadratic trends were highly significant (*p* < 0.0001) for the F1 amplitudes in all conditions. These results suggest that the FFR F1 also showed categorical coding regardless of attention or presentation order.

We measured the F0 of the first and last tokens in each train to determine if neural adaptation occurred given the rapid, clustered nature of our stimulus presentation. We found no difference in F0 amplitude between the first and last response in each train (*F_1,647_* = 0.20, *p* = 0.6551; data not shown). This confirms there was little to no adaptation of brainstem responses due to the rapid succession of auditory stimuli (Bidelman and Powers, 2018) and thus rules the confound that serial order effects in the FFR data were due to mere neuronal fatigue.

**Figure 8** shows response-to-response correlations between Tk4 (ambiguous token) and Tk1/7 (prototypical tokens) FFRs as a function of presentation order. We found a main effect of presentation order (*F_1,45_* = 4.39, *p* = 0.0417) and a significant interaction between presentation order and token (*F_1,45_* = 4.92, *p* = 0.0317). The interaction suggests Tk4 FFRs showed stronger similarity to Tk1 in the forward direction but stronger correspondence to Tk7 in the backward direction. By token, the direction contrast was stronger at Tk1 (*p*=0.0038) than Tk 7 (*p*=0.93). These findings suggest that the FFR to an otherwise identical (and categorically ambiguous) speech token was modulated by perceptual state. That is, FFR neural representations were warped toward the direction of the vowel prototype under each stimulus context (i.e., mirroring Tk1 for forward stimulus ordering and Tk7 for backward stimulus ordering).

**Figure 8.**
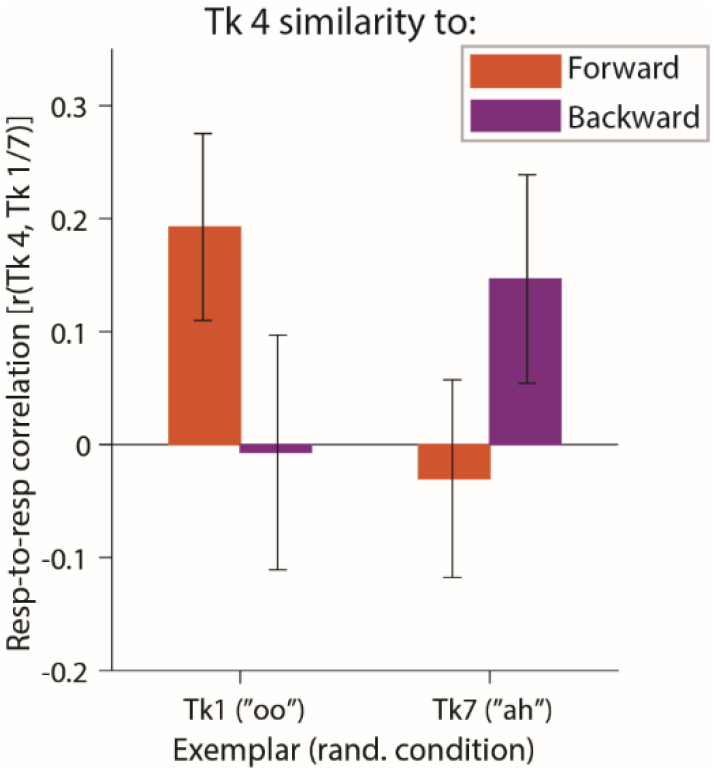
FFRs show category-specific coding. Comparison of response-to-response correlations between FFRs to the ambiguous speech token (Tk4) presented in backward and forward conditions with responses to either prototypical vowel (Tk1/7). Higher correlation coefficients indicate a stronger similarity to that speech category (i.e., /u/ or / /). Errorbars = =±1 s.e.m.

Neural classifier performance is shown in **Figure 9**. Single-trial decoding was expectedly robust for classifying the phoneme endpoints of the continuum (i.e., Tk1 vs. Tk7). At the group level, cross-validated accuracy was 86% AUC (d-prime=1.49), resulting in few confusions between true and predicted token labels and thus highly discriminable responses (Χ^*2*^ = 8468, *p*<0.00001). Individual vowel decoding from FFRs was equally good and well above chance levels (one-sample t-test against 50%: *t_14_* = 27.62, *p*<0.0001). These control decoding analyses indicate that phonetic properties of vowel prototypes (i.e., /u/ vs. / /) were easily distinguished via spectrotemporal features carried in FFRs.

**Figure 9.**
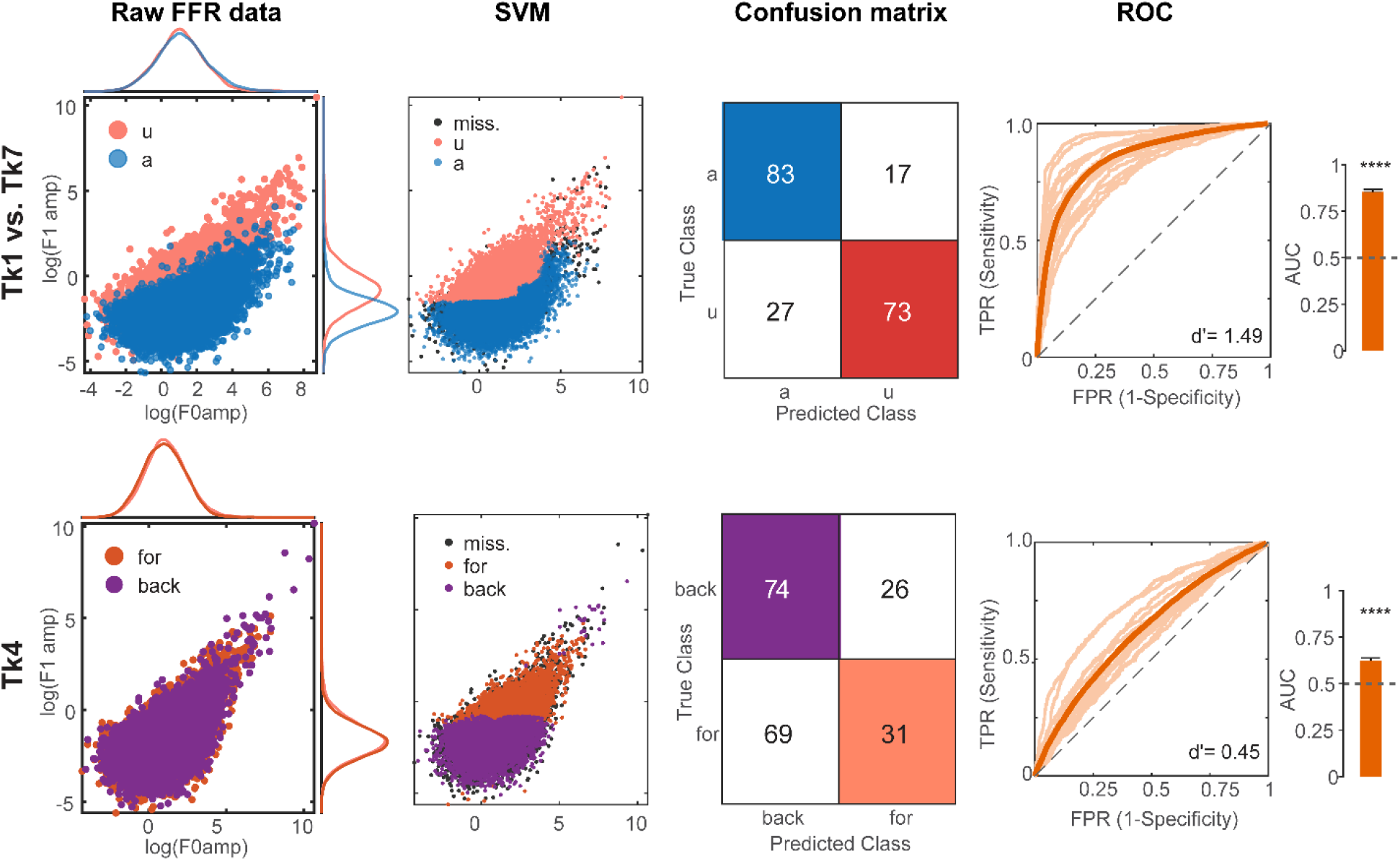
Single trial FFR decoding reveals evidence of phonetic encoding and perceptual warping depending on stimulus presentation order. SVM classification of FFRs decoding (*top row*) category prototypes (Tk1=/ / vs. Tk7=/u/) and (*bottom row*) stimulus order (direction) for ambiguous Tk4 responses. *First column*, raw FFR F0 and F1 amplitudes extracted from N=27000 single trial FFRs. *Second column*, SVM output showing the decision boundary and posterior class labels predicted after SVM training. Black=misclassified observations. *Third column,* cross-validated confusion matrices show the proportion of predicted vs. true class labels. Higher values along the diagonal denote more successful separability of FFRs and better decoding performance. *Fourth column*, ROC curves. Thin lines= single subjects; thick line; grand average across participants; dotted line= chance performance. *Fifth column,* average ROC area under the curve (AUC) across listeners for classifier performance (cf. %-accuracy). (*Top row*) Prototypical categories (Tk1 vs. Tk 7) are easily distinguished via neural FFRs, resulting in few confusions and high classification accuracy. Note the large separability in F1 (but not F0) amplitude measures in the raw data. (*Bottom row*) Decoding FFRs to Tk4 presented in the forward vs. backward serial directions, which produce perceptual hysteresis. Decoding performance is well above chance even in light of the low separability of the data (i.e., all Tk4 trials) suggesting FFRs contain adequate information to code listeners’ trial-by-trial speech percepts. TPR, true positive rate. FPR, false positive rate. AUC, area under the (ROC) curve. \*\*\*\**p*<0.0001; Errorbars = ±1 s.e.m.

Having confirmed FFRs carry sufficient information on the category identity of speech signals, we next asked whether responses to category ambiguous speech (Tk 4) showed differential encoding depending on the direction of stimulus presentation. As confirmed behaviorally, forward vs. backward serial ordering of the stimulus continuum induces perceptual hysteresis that changes listeners’ percept of otherwise identical speech sounds (see Fig. 2). Single-trial decoding was expectedly poorer for this more challenging classification problem given lower separability of the data. However, FFRs were surprisingly distinguishable based on stimulus order (i.e., Tk4_for_ vs. Tk4_back_). Group level classification accuracy was 62% AUC (d-prime=0.45) with more frequent vowel confusions—the SVM tended to predict more Tk4 responses as stemming from the backward condition, suggesting a slight bias in labeling. Still, the confusion pattern was highly discriminable (Χ^2^ = 66.5, *p*<0.00001) and individual decoding was well above chance (*t*_14_ = 9.26, *p*<0.0001). These neural decoding results complement the response-to-response correlations and indicate that FFRs to otherwise identical speech stimuli are warped on a trial-by-trial basis according to the surrounding stimulus context. In subsequent analyses, we focus on the relevance of these dynamic neural effects to *behavioral* categorization of speech.

### 3.3 Brain-behavior relationships

We used GLME regression models to determine whether neural FFR measures predicted aspects of listeners’ categorical perception. For β0 (boundary location), the multivariate model indicated FFR measures predicted ∼63% of the variance in behavior (*R^2^* = 0.63). Evaluating individual terms revealed a significant predictor in FFR F1 frequency on listener’s categorical boundaries (*t_11_*= 2.43, *p* = 0.03). For β1 (psychometric slopes), the multivariate model predicted ∼71% of the variance (*R^2^*= 0.71). Evaluating individual terms revealed a significant predictor in FFR F0 frequency on listener’s psychometric slopes (*t_11_* = 2.87, *p* = 0.015). These results suggest the degree to which subcortical responses code different speech features predict properties of listeners’ vowel categorization.

## 4. DISCUSSION

The current study measured brainstem FFRs concurrent with behavioral responses to acoustic-phonetic continua presented in various stimulus orderings (sequential vs. random presentation) and attentional states (active vs. passive tasks). Our innovative stimulus task establishes a new paradigm to obtain FFRs and behavioral responses to speech concurrently. Using this novel approach, we show that attention modulates the encoding of speech as early as the auditory midbrain and moreover, that FFRs encode speech categorically.

### FFR responses obtained concurrent with active task

Most speech-FFR studies drawing putative links between auditory brainstem coding and aspects of speech perception have used passive listening tasks (Aiken and Picton, 2008; Bidelman et al., 2013a; Skoe and Kraus, 2010; Slugocki et al., 2017). This has led to claims that FFRs reflect a perceptual correlate of behavior. However, in the absence of an active perceptual task in previous work, establishing this link is spurious. Recent advancements in stimulus paradigms have shown that active, perceptual challenging tasks can induce modulations in the speech-FFR, revealing brainstem representations are subject to attentional gain modulation (Price and Bidelman, 2021). Through use of our innovative clustered stimulus paradigm, we further demonstrate a feasible method to obtain speech-FFRs simultaneous with an active behavioral speech listening task. Our data provide new and important evidence that speech-evoked brainstem responses, likely their cortical ERP counterparts (Carter et al., 2022), are actively modulated by listeners trial-by-trial perception of the speech signal and its surrounding context. Consequently, we infer FFRs reflect more than mere sensory-acoustic representations, and instead carry true perceptual correlates of the speech signal.

Behaviorally, we found that the slopes of listeners’ psychometric functions were steeper in sequential vs. random presentation ordering. This agrees with previous findings (Carter et al., 2022) and suggests that sequential presentation solidifies categorization as individuals rapidly decide what category the sound associates with. Additionally, the serial presentation of tokens in our paradigm likely strengthens the sensory (echoic) memory trace which would reinforce individuals’ decision by the time they execute their behavioral response (Näätänen et al., 2007; Winkler et al., 1993).

Surprisingly, RTs were slower in backward vs. both the forward and random conditions. On the contrary, we would have expected the random condition to produce longer RTs than either sequential condition. RTs may have been slower in the backward condition due to a greater salience of rising than falling frequency stimuli (Carter et al., 2022; Luo et al., 2007; Schouten, 1985). Perhaps in our paradigm, listeners subtly slowed their identification to ensure they were selecting the correct sound, whereas in forward and random conditions, the change in F1 frequency was perceptually salient enough to keep RTs rapid. Additionally, RT patterns in conventional speech categorization tasks typically show an inverted U shape across the continuum, with RTs slowing around the categorical boundary compared to the prototypical tokens (Pisoni and Tash, 1974). We did not observe this in the current study. This may relate to listeners deciding their percept early in the stimulus train, then selecting their response once the train ends. That is, RTs might be locked more to the ending of the entire stimulus train than to the processing of the phoneme, *per se*. Despite the lack of token effect, RTs were however modulated by the overall ordering of speech, indicating that decision speeds can be facilitated by recent stimulus history (i.e., context).

### Brainstem FFRs are modulated by attention

Strikingly, we found that speech-FFRs (F0 amplitudes) were much larger in active vs passive conditions, confirming that attention actively shapes neural encoding at the brainstem level. We had expected to also see changes in FFR F1 amplitudes as a function of presentation order, but this was not observed (see **Fig. 7**). Attention effects in the FFR thus seem localized to low-frequency components of the speech signal (Holmes et al., 2018). The effect of attention on any property of brainstem responses has been equivocal in previous work; some studies support (Galbraith et al., 1998; Hartmann and Weisz, 2019; Price and Bidelman, 2021) and others refute (Aiken and Picton, 2008; Dunlop et al., 1965; Galbraith and Kane, 1993; Varghese et al., 2015) attentional effects on FFRs.

Attentional modulation of FFRs observed here is presumably driven by corticofugal fibers that enhance brainstem activity selectively according to perceptually-relevant information in cortex. Animal studies have shown the corticofugal fibers shape subcortical function during short-term learning (Bajo et al., 2010; Suga, 2008). In humans, corticofugal mechanisms are thought to be particularly important in difficult speech-listening environments (Lai et al., 2022a; Price and Bidelman, 2021). These effects could relate to the short-term memory modulation caused in nonlinear dynamical processing of speech, wherein the encoding of ambiguous speech tokens at lower levels are continuously shaped by higher cortical structures. Indeed, perceptual warping effects on primary auditory cortex responses are thought to arise from prefrontal memory areas (Carter et al., 2022). It is possible such perceptually-relevant biasing percolates back to even more peripheral auditory areas (i.e., brainstem) as suggested by the category tuning of FFRs observed here. In this regard, corticofugal fibers might carry category identify from cortex further down the system, rending changes in speech representations at the brainstem level. Previous anatomical work has demonstrated cortico-collicular connections originating in the frontal lobes and terminating in the brainstem that contain GABAergic and glutamatergic neurons. These connections are thought to shape responses in inferior colliculus, a major source of the FFR, through complex excitatory and inhibitory interactions (Olthof et al., 2019). Consequently, the necessary circuitry is in place for higher-level brain regions (frontal lobe) to modulate early signal encoding in the FFR (see Carter et al., 2022 for similar evidence between frontral lobe and auditory cortex). Our data provide strong evidence of attentional modulation of subcortical responses, possibly originating in the distal frontal lobes. Though future studies are needed to confirm this hypothesis.

Our task requires listeners to perform online categorization judgments and continuously monitor the speech stimuli. Previous tasks evaluating brainstem-attention effects have used simple tasks (e.g., counting, detection, attention redirection, etc.) (Galbraith et al., 1998; Galbraith and Kane, 1993; Varghese et al., 2015) or oddball paradigms (Hartmann and Weisz, 2019; Price and Bidelman, 2021), which may allow listeners to periodically disengage from the task and fail to produce FFR-attention effects. We have recently shown that task disengagement has strong influences on cortical arousal which simultaneously causes fluctuations in speech FFR responses (Lai et al., 2022a). Our task arguably requires more sustained attention which may account for the much larger (x2-3) brainstem attentional effects we find in the present study compared to previous reports (Price and Bidelman, 2021).

### Brainstem FFRs carry category-level information (perceptual correlates) of speech

Another novel finding revealed by our innovative task is that FFRs encode speech in a categorical fashion. Category representations in the FFR is unlikely to be *de novo* (i.e., local) to the midbrain. Rather, we posit that corticofugal fibers modulate early sound encoding of the stimulus to fit the perception of the token (Suga, 2008; Suga et al., 2000). We have previously shown that at the level of cortex (Carter et al., 2022), activity in frontal brain regions influences the encoding and subsequent perception of category-ambiguous speech sounds (cf. Tk4) (Carter et al., 2022). Previous work has also demonstrated that changes in perception can drive enhancements of the FFR (Cheng et al., 2021), suggesting speech processing is influenced by predictions of the percept. Our response-to-response correlations and neural decoding results support perceptual encoding in the FFR. Brainstem responses to otherwise ambiguous speech tokens were biased towards a given prototype depending on the direction of presentation. These results were independent of neural adaptation ruling out explanations that our FFR warping effects where driven by the normal physiological byproducts of rapid auditory processing (i.e., refractory of neuronal firing). Together, this indicates the FFR is not merely a passive representation of the acoustic speech signal but is dynamically shaped by higher-order perceptual processes, and by surrounding stimulus context. Such active modulation of brainstem representations might help simplify speech decisions upon arrival to auditory cortex (Asilador and Llano, 2021; Lesicko and Geffen, 2022).

Importantly, our speech stimuli were designed with F0s well above cortical phase locking limits as reported in both humans and animal models (Brugge et al., 2009; Gnanateja et al., 2021; Guo et al., 2021; Joris et al., 2004; Wallace et al., 2000). This rules out the possibility that our FFR results are conflated by cortical contributions (Coffey et al., 2016b). This, in addition to the lack of neural adaption (characteristic of more peripheral auditory nuclei), strongly supports a brainstem locus of our findings.

The F0 amplitudes in neural responses to / / stimuli were larger than in responses to /u/ stimuli, unexpectedly revealing differences due to perception. These differences could be due to an overall offset of the root mean square amplitude for the response being demonstrated in the F0. Alternatively, changes in F1 have previously been revealed to increase F0 amplitude when F1 contains linguistically-relevant cues (Krishnan et al., 2011). We can rule out explanations due to loudness differences, as all tokens were matched in sound level and perceptual loudness (94.1±1.0 phon) (Moore et al., 1997). This suggests that acoustic differences do not underlie the differences we find in FFR F0 amplitude, but instead indicate that perceptual differences between sides of the continuum drive this neural differentiation. This difference in FFR F0 also cannot be attributable to acoustical F0 differences. In fact, an acoustic analysis showed the reverse pattern as in the FFR; stimulus F0 of the /u/ tokens was 250% higher than amplitudes of / / tokens. Meanwhile, the FFR F0 amplitude to /u/ tokens were smaller, i.e., 47.9% the amplitude as / / tokens. The dissociation of acoustics and neural responses indicates a neural, rather than acoustic, origin for the differences we observed. This view that perception drives FFR differences is further bolstered by the similarities between neural responses to tokens within the same phonetic category.

Two alternative theories suggest how ambiguous phonemes are categorized and might account category-level coding we find in FFRs: the Natural Referent Vowel (NRV) and the Native Language Magnet (NLM) models. The NRV proposes that spectral prominences that are easier to detect lead to directional asymmetries in category discrimination tasks. Contrastively, the NLM proposes that directional asymmetries are caused by the vowel space being biased towards native phonetic prototypes—built through long-term statistical learning— which act as perceptual magnets for ambiguous phonemes (Masapollo et al., 2017; Zhao et al., 2019). Our findings have support from both models, but more strongly support the NLM within the context of categorization. Perception of ambiguous tokens was driven by listeners categorizing the sound as one of the prototypical tokens (i.e., NLM). This interpretation is further bolstered by the U shape found in FFR responses, which suggests there is pull from both prototypes. However, the fact Tk4 responses were more strongly correlated with Tk1_FOR_ than Tk7_BACK_ responses suggests there may be a slight bias towards vowels with more prominent F0/F1 configuration, consistent with NRV. It could be that the vowel biasing is more strongly related to NLM in our study as we used a continuum with prototypes that were both native to our listeners. Previous studies supporting NRV interpretations of FFR and direction-dependent vowel coding effects compared within category stimuli (Masapollo et al., 2017; Zhao et al., 2019).

Additional evidence that the FFR reflects aspects of speech percepts was our finding that response components were associated listeners’ categorical boundary and the slope of their psychometric function. We and others have shown that perception begins to differentiate phonemes categorially early in the cortical hierarchy and no later than primary auditory cortex (Bidelman and Lee, 2015; Bidelman and Walker, 2019; Carter and Bidelman, 2021; Chang et al., 2010). Here, we extend these findings by showing category-specific neural representations extend as low as the brainstem. As the FFR is largely driven by midbrain regions (Bidelman, 2018b), the link between FFR and psychometric measures is consistent with notions that low-level auditory representations carry information regarding signal clarity, strength of categorization, and vowel identity (Binder et al., 2004). In contrast, FFR measures did not predict the speed of listeners’ decisions. RTs are however largely driven by higher-order frontal brain regions (Binder et al., 2004), so it is perhaps not surprising that FFRs failed to predict perceptual speeds (but see Galbraith et al., 2000). Collectively, the fact that FFRs are strongly modulated by attention and show category-specificity (**Fig. 7**) strongly suggests brainstem FFRs carry perceptual correlates related to how listeners ultimately hear the speech signal.

Broadly, our findings indicate top-down processes modulate brainstem representations to fit the anticipated speech percept. Our data bolster notions that the FFR carries perceptually relevant cues related to phonetic representations and is thus more than just a neural mirror of the acoustic signal. Together, our findings suggest that midbrain plays a vital role in the active perception and categorization of speech.

## Acknowledgements

Work supported by the National Institute on Deafness and Other Communication Disorders (R01DC016267). Requests for data and materials should be directed to G.M.B.

